# Computational investigation unveils pathogenic LIG3 non-synonymous mutations and therapeutic targets in acute myeloid leukemia

**DOI:** 10.1101/2025.02.22.639676

**Authors:** Md. Arif Hossen, Umme Mim Sad Jahan, Md. Arju Hossain, Khalid Hossain Asif, Ahona Rahman, Sabbir Ahmed, Md. Moin Uddin, Md Faisal Amin, Muhammad Abdul Bari, Mohammod Johirul Islam, Mohammad Kamruzzaman, Soharth Hasnat, Mohammad Nasir Uddin, Tofazzal Islam, M. Nazmul Hoque

**Author notes:** **Correspondence to:** Mohammad Nasir Uddin, Tofazzal Islam, and M. Nazmul Hoque.

## Abstract

Single nucleotide polymorphisms (SNPs) in DNA repair genes can impair protein structure and function, contributing to disease development, including cancer. Non-synonymous SNPs (nsSNPs) in the *LIG3* gene are linked to genomic instability and increased cancer risk, particularly acute myeloid leukemia (AML). This study aims to identify the most deleterious nsSNPs in the *LIG3* gene and potential therapeutic targets for DNA repair restoration in AML. We employed in-silico computational methods to analyze *LIG3* nsSNPs, using PredictSNP and Mutation3D to assess pathogenicity. Subsequently, molecular docking and dynamics simulations were conducted to evaluate ligand-binding affinities and protein stability. Out of the 12,191 mapped SNPs, 132 were nsSNPs located in the coding region. Among these, 18 nsSNPs were identified as detrimental including 12 destabilizing and 6 stabilizing nsSNPs. Nine cancer-associated nsSNPs, including L381R and R528C, were predicted due to their structural and functional impacts. Further analysis revealed key phosphorylation and methylation sites, such as 529S and 224R. Molecular dynamics simulations highlighted stable interactions of compounds AHP-MPC and DM-BFC with wild-type and R528C mutant LIG3 proteins, while R671G and V781M mutants showed instability. Protein-protein interaction networks and functional enrichment linked LIG3 to DNA repair pathways. Kaplan-Meier analysis associated high *LIG3* expression with improved survival in breast cancer and AML, suggesting its role as a prognostic biomarker. This study emphasizes the mutation-specific effects of *LIG3* nsSNPs on protein stability and ligand interactions. We recommend identifying DM-BFC to advance personalized medicine approaches for targeting deleterious variants, following *in vitro* and *in vivo* validation for AML treatment.

## 1. Introduction

Acute Myeloid Leukemia (AML) is a prevalent and deadly leukemia characterized by the swift expansion of myeloid progenitor cells in the bone marrow, leading to immense disruption of normal hematopoiesis and representing a fatal form of bone marrow malignancy [1, 2]. AML affects 12.6 out of 100,000 persons in the US who are 65 years of age or older at its peak, with an annual incidence of around 2.4 per 100,000. The incidence rises steadily with advancing years [3]. DNA repair is a crucial molecular defense system against chemicals that cause cancer, degenerative diseases, and aging. Various repair systems exist in humans to defend the genome by fixing changed bases, DNA adducts, cross-linkages, and double-strand breaks (DSBs) [4]. DSB repair is crucial for maintaining genomic integrity and preventing mutations that can lead to cancer and other diseases. In higher eukaryotes, DNA DSBs are predominantly repaired using a simple mechanism called non-homologous end joining (NHEJ). NHEJ involves the ligation of broken ends without the requirement of homology [5]. Translocations of chromosomes are facilitated by alternative non-homologous end-joining (alt-NHEJ), which is a newly discovered process for repairing DNA DSBs [6]. All kinds of leukemia exhibit impaired DSB repair, while several key components of DSB repair are especially affected. The Ku70/80 complex and DNA-dependent protein kinase (DNA-PK) play a role in the non-homologous end-joining process [7].

LIG3, encoding DNA Ligase III, is crucial in the emergence of AML due to its role in DNA repair mechanisms, especially in NHEJ. This pathway frequently demonstrating heightened expression in cancers indicated by genomic instability, like AML, in which deficiencies in DNA repair serve a vital part in cancer progression and resistance to treatment [8, 9]. LIG3 becomes more active when the normal system, which depends on DNA ligase IV, is not functioning properly [10]. LIG3 can be divided into two forms, LIG3-α and LIG3-β, using various splicing procedures. LIG3-α participates in the repair of nucleic acids through the DNA repair protein XRCC1, whereas LIG3-β is found in male germ cells [11]. *LIG3* is necessary for the metabolism of mitochondrial DNA. *LIG3* interacts with the single-strand break repair protein XRCC1 through its C-terminal BRCT domain. *LIG3* has been shown to possess end-joining activity in cellular extracts and in *LIG4*-deficient cells that were depleted of *LIG3* using plasmid substrates. This suggests that *LIG3* is involved in a secondary mechanism of NHEJ for repairing DSBs [12].

SNPs denote variations in DNA sequences resulting from a mutation of a single nucleotide at the genomic level. The human genome is estimated to encompass a minimum of 3 million SNPs, with an average frequency of occurrence every single 300 base pairs [13]. SNP technologies are valuable for studying differences in treatment responsiveness between individuals and finding genes that cause human diseases. Moreover, the biological mechanisms behind sequence evolution can be understood by utilizing SNPs [14]. SNPs play a crucial role as markers in numerous research that establish connections between variations in DNA sequences and changes in observable traits [15]. nsSNPs and mutations have been associated with human features and diseases [16]. SNPs can also impact gene expression and protein function and are observed throughout numerous genomic locations, including as promoters, exons, and introns. Finding SNPs may facilitate the disease severity anticipation and personalized therapeutic strategies. The expression levels of LIG3 are associated with prognosis in several cancers, suggesting that both SNPs and expression may function as markers for clinical results. Particularly LIG3 SNPs have been linked to somatic mutations affecting numerous malignancies, underscoring their likelihood of prognostic significance [17]. *LIG3* gene can facilitate NHEJ even when *LIG4* gene is not present, as well as nucleotide excision repair (NER) and homologous recombination repair (HRR) [18]. Gene polymorphisms associated with DNA repair pathways, such as *LIG3*, might contribute to the initiation and progression of Alzheimer’s disease [19]. The LIG3-XRCC1 pathway identifies ADP-ribosylation and is necessary for the joining of Okazaki fragments in the final stages of DNA replication [18]. A number of studies have examined mutations in the *LIG3* gene; however, the prediction of harmful SNPs in the *LIG3* gene linked to AML has not yet been performed. Further investigation is required to figure out the most deleterious and disease-associated SNPs in the *LIG3* gene associated with other malignancies, including AML. Our computational analysis points out that the R528C, R671G, and V781M mutations in the LIG3 gene significantly affect protein structure and function. These insights might shed light on why some mutations are associated with an increased risk of disease. Therefore, our objective was to identify the most deleterious nsSNPs in the *LIG3* gene and potential therapeutic targets for DNA repair restoration in AML. Additionally, we identified therapeutic targets with potential to mitigate the effects of these mutations while improving protein composition, stability, and function, opening new possibilities for cancer treatment.

## 2. Materials and Methods

**2.1 Retrieval of LIG3 nsSNPs dataset**

The dataset of SNPs for the human *LIG3* gene and its protein sequence (Uniprot ID: P49916) was obtained from the NCBI dbSNP (https://www.ncbi.nlm.nih.gov/snp/) (Accessed on: 5 May, 2024) and UniProtKB (https://www.uniprot.org/) (Accessed on: 5 May, 2024) databases, respectively. A total of 12,191 SNPs that belong to different functional classifications (**Fig. 1**) were mapped within the *LIG3* gene sequence. Among 12,191 SNPs, 132 were non-synonymous SNPs (nsSNPs) located in the coding area, which may result in missense or nonsense mutations, hence influencing the structure and function of the protein. In this study, our focus was on the coding region of the LIG3 protein, where we evaluated the nsSNPs. Subsequently, the 132 identified nsSNPs were extracted and subjected to detailed analysis.

**Fig. 1.**
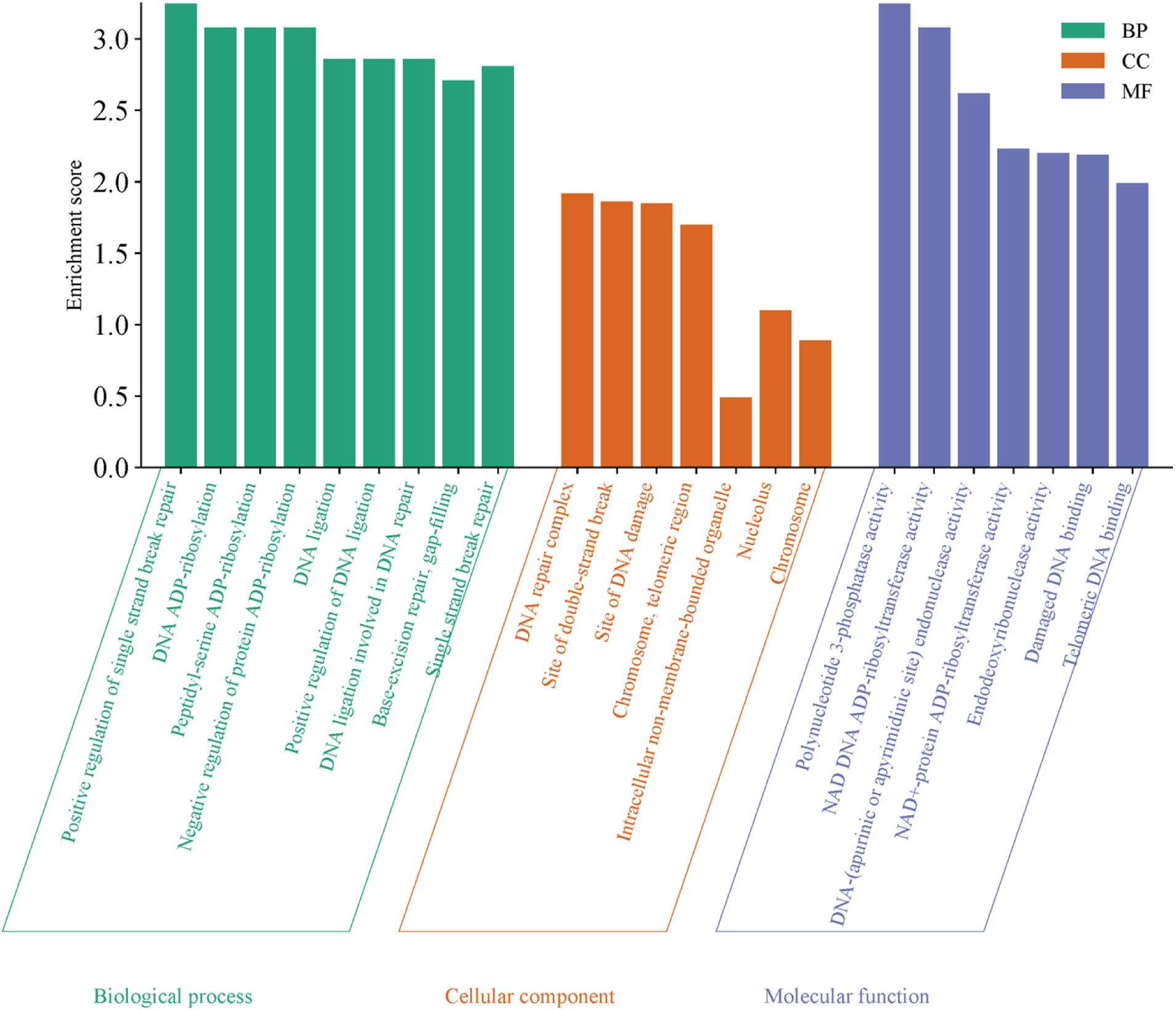
Assessment of the LIG3 gene with a deep focus on Gene Ontology (GO) pathways, specifically Biological Process (BP), Cellular Component (CC), and Molecular Function (MF).

### 2.2 Screening the highly deleterious nsSNPs

We used different bioinformatics tools to assess nsSNP variations, thoroughly screening and prioritizing alterations predicted to have detrimental effects. To predict the consequences of detrimental SNPs in the human genome, PhD-SNP (https://snps.biofold.org/phd-snp/phd-snp.html) (Accessed on 10 May, 2024) tools was utilized which uses Support Vector Machines (SVMs) as classifiers and seems for alterations in protein sequence, mutation locations, and mutated residues [20]. PANTHER is a biological and evolutionary database for all genes that code for proteins was utilized for categorizing the genes based on the evolutionary trajectory and functional attributes [21]. Using the availability or frequency of protein substitutions in the sequence of query protein, PolyPhen-2 categorizes mutations as potentially lethal (>0.15), likely detrimental (>0.85), or benign, depending on their impact on protein expression [22, 23]. Subsequently, Predict-SNP, a consensus algorithm that incorporates the MAPP, SNAP, and PolyPhen-1, provides data for each mutation and significantly enhances estimated performance, proving that consensus prediction is a trustworthy and precise alternative to predictions generated through distinct tools [24]. Mutated protein function was examined using SIFT to ascertain if nsSNPs had a positive or negative effect [25]. Integrated computational methods, including functional annotation of SNPs through conservation profiling, analysis of protein structural and functional data, and linkage of coding SNPs to gene transcripts, enable a comprehensive evaluation of the likelihood of harmful missense mutations.

### 2.3 Functional consequences of nsSNPs on protein

We evaluated protein stability using the SVM-based web server I-Mutant 2.0 (Accessed on May 30, 2024). This method was pivotal to our study due to its capability to predict stability changes resulting from mutations [26]. MUpro (Accessed on May 31, 2024) was employed to analyze changes in protein sequences by comparing residues between the wild-type and mutant proteins [27]. Additionally, we utilized the ΔΔG free energy change values to assess protein stability through the mutation cutoff scanning matrix (mCSM) method (Accessed on May 31, 2024) [28]. A ΔΔG value greater than 0 signifies enhanced protein stability, while a value below 0 indicates that the mutation adversely affects protein function [29, 30].

### 2.4 Assessment of nsSNPs linked to cancer

We analyzed a set of amino acid substitutions arising from somatic cancer mutations using Mutation 3D (http://www.mutation3d.org/) (accessed on June 3, 2024). This tool is widely utilized to evaluate the effects of cancer-associated nsSNPs on protein function and disease progression. By employing a 3D clustering approach, the tool identifies potential cancer-driving alterations in a protein’s amino acid composition. The analysis required input data comprising the target protein and its associated mutations [31].

### 2.5 Structural and functional changes prediction

MutPred2 (accessed on June 7, 2024) was utilized as a method for predicting structural and functional alterations induced by amino acid variants [32]. This tool enhances the detection of harmful variants by simulating how mutations impact protein structure and function, aiding in the understanding of disease mechanisms. It also provides insights into the specific biological pathways involved in disease progression. Protein FASTA sequences, along with amino acid variations, were input into MutPred2 for analysis. The method emphasizes the integration of genetic and molecular data through machine learning techniques [33].

### 2.6 Anticipating the modifications of protein 3D structure resulting from mutation

To assess the impact of residue substitutions on protein structure, we utilized the Project Hope server (https://www3.cmbi.umcn.nl/hope/) (accessed on June 15, 2024). This tool integrates 3D structural data and provides detailed insights into the structural differences between native and mutant protein residues. Structural impact analysis was performed using the protein sequence of LIG3 and its mutations (nsSNPs) [34]. We examined how alterations in amino acid composition affect native structures, focusing on differences in hydrophobicity, charge, and size between wild-type and mutant residues. Understanding these modifications’ influence on the protein’s three-dimensional structure is vital for elucidating its function, guiding future experimental studies, and developing novel treatments and diagnostic tools [35].

### 2.7 Estimation of post translation modification sites

Several computational tools, such as NetPhos 3.1

(https://services.healthtech.dtu.dk/services/NetPhos-3.1/) (accessed on June 20, 2024) and GPS-MSP 1.0 (http://msp.biocuckoo.org/), have made significant advances in identifying post-translational modification (PTM) sites, particularly for phosphorylation and methylation. These tools use machine learning and deep learning techniques to enhance prediction accuracy by incorporating sequence and structural data. We employed the NetPhos 3.1 tool, which utilizes multiple neural networks to predict phosphorylation sites on tyrosine, threonine, and serine residues, identifying potential locations for these modifications [36]. In addition, we used GPS-MSP 1.0 to predict potential methylation sites along the protein chain [37].

### 2.8 Protein-protein interaction network prediction

Protein-protein interactions (PPIs) were analyzed using the STRING database (v12.0) (accessed on June 22, 2024) [38]. Interaction networks were constructed based on experimental data, co-expression patterns, curated databases, and text mining. A confidence score threshold of ≥0.7 was applied to ensure high-confidence interactions. The generated networks were visualized and exported for further analysis. Functional enrichment and clustering tools within STRING were employed to identify key pathways and interaction modules [38]. Analyzing the PPI data is essential, as mutant proteins can continuously influence other proteins in the diseased state. Understanding these interactions provides insights into the underlying mechanisms of clinical conditions and aids in identifying the source protein and its associated network [38, 39].

### 2.9 Exploring pathways with gene ontology and KEGG enrichment

Gene Ontology (GO) is a widely used knowledge-based resource that provides organized and computable information on gene functions, focusing on biological processes (BP), cellular components (CC), and molecular functions (MF). These three ontologies are integral to GO enrichment analysis [40]. We performed GO analysis using EnrichR (https://maayanlab.cloud/enrichr-kg) (accessed on June 30, 2024) to identify statistically significant associations (P < 0.05) between the input gene set and curated databases covering CC, MF, and BP [40]. Following this, we utilized SRplot (accessed on June 30, 2024) to create graphical summaries that visualize the enriched analysis results [41].

### 2.10 Superimposition and molecular layering of wild-type and mutant-type proteins

Protein structure superimposition is a key technique in structural biology for comparing protein structures to understand their evolutionary relationships, functions, and dynamics. By aligning protein structures, it reveals similarities, differences, and dynamic changes over time, highlighting functional patterns [42]. Chimera 1.16 was used to superimpose the native LIG3 protein and its mutant variants [43].

### 2.11 Molecular docking, pharmacokinetic and dynamics simulation profiling

Molecular docking is a key technique in drug discovery, facilitating virtual screening and drug repurposing [44]. Using Autodock Vina (v1.2.1), we evaluated how detrimental mutations affected LIG3’s (UniProt ID: P49916) binding affinity [45]. The LIG3 crystal structure complex, obtained from RCSB (https://www.rcsb.org/; Accessed on July 10, 2024 ) and analyzed with Phyre2 [46], was energy-minimized using Swiss-Pdb Viewer [47], to generate mutant forms. We focused on four proteins: the wild-type and three mutant LIG3 variants with nsSNPs. Sixteen ligands from PubChem (CIDs: 707801, 70687578, 116535, 59937, 408383, 676443, 718154, 722325, 609964, and 684700) were docked with both wild-type and mutant proteins [48]. Ligands were converted to pdbqt format using Autodock Vina, and grid boxes (x = 77.5488082123; y = 72.4516607666; z = 73.5725282574) were optimized for docking efficiency. Docking results and ligand-protein interactions were visualized using BIOVIA Discovery Studio (v21.1.0) [49].

The pharmacokinetic phase (absorption, distribution, metabolism, excretion) and along with toxicity study (ADMET) are some essential parameters in designing and development of new drug. An in-silico computational pharmacokinetics approach was used to determine the ADMET properties of the AHP-MPC (CID: 70687578) and DM-BFC (CID: 707801). In cases of AML, breast cancer, hepatocellular carcinoma, and other diseases, these compounds may be evaluated as possible therapeutic agents that target LIG3 [48]. Drug-likeness and pharmacokinetics parameters such as Absorption, Distribution, Metabolism, and Excretion (ADME) in the compounds were evaluated through the SwissADME [50] and pkCSM [51] web tools. For toxicity prediction, we used Protox III online server [52]. In this analysis, the Simplified Molecular Input Line Entry System (SMILES) formats of the both compounds were retrieved from the PubChem database. Lipinski’s rule of five was used to assess the drug-likeliness properties of the compounds [53]. Molecular dynamics simulations (MDS) were performed using Desmond v24, developed by Schrödinger LLC, to validate the interactions predicted during docking analysis [54]. MDS applies Newton’s classical laws of motion to compute atomic positions and velocities over time, generating new configurations at small intervals. This approach enables the prediction of ligand-binding behavior under near-physiological conditions, providing dynamic insights into the stability and interaction patterns of the ligand-protein complexes [55]. The ligand-receptor complexes were preprocessed using the Protein Preparation Wizard, which facilitated optimization, energy minimization, and the addition of any missing residues to the protein complexes. The System Builder application was subsequently employed to construct the simulation system. The TIP3P solvent model, featuring an orthorhombic box structure, was used along with the OPLS_2005 force field, under conditions of 300 K temperature and 1 atm pressure, to ensure a realistic simulation environment [56, 57]. Each complex was neutralized using counter ions and 0.15 M sodium chloride to replicate physiological conditions. The simulation trajectories were recorded at 100 ps intervals and visualize with module of simulation interaction diagram

### 2.12 Clinical validation of LIG3

To assess the prognostic significance of LIG3 gene expression, survival analysis was performed using the Kaplan-Meier Plotter, an established online tool that integrates clinical and gene expression data across various cancer types [58]. Kaplan-Meier survival curves were generated to evaluate the relationship between LIG3 mRNA expression levels (high vs. low) and overall survival (OS). Hazard ratios (HR) with 95% confidence intervals (CI) and log-rank P-values were calculated to determine statistical significance. Patients were stratified into high- and low-expression groups based on median expression levels. A statistical threshold of P < 0.05 was applied, highlighting the relevance of the findings.

## 3. Results

### 3.1. Assessment of deleterious nsSNPs

We retrieved 12,191 nsSNPs within the *LIG3* gene from dbSNP. Of these, 902 (7.4%) were missense variations, 9,685 (79.44%) were intronic variants, 398 (3.26%) were synonymous variants, 132 (1.08%) were somatic missense variants, and 1,074 (8.81%) belonged to other categories (**Fig. S1**). The missense variations underwent further analysis to pinpoint the most harmful SNPs. Notably, 132 nsSNPs were identified and categorized as somatic (**Table S1**). Among these, we identified 18 detrimental nsSNPs that potentially affect the overall structure or function of the LIG3 protein (**Table S2**).

### 3.2 Prediction the effects of nsSNPs and post-translation modification on protein stability

To evaluate protein stability, we examined the 18 detrimental nsSNPs and found that 12 of them significantly reduced protein stability, while the other variants improved it. These findings were supported by reliability index (RI) values and ΔG free energy change values (**Table S3**). A decrease in stability indicates protein destabilization, while an increase suggests stabilization. For further analysis, we concentrated solely on the missense variants of nsSNPs. Using the GPS-MSP 1.0 tool, we identified 224R as a potential methylation site on the LIG3 protein. Additionally, phosphorylation site predictions from NetPhos 3.1 identified 529S in the native protein and 666Y in the mutant as potential phosphorylation sites (**Fig. S2, Table S4**).

### 3.3 Evaluation of cancer-associated nsSNPs and their structural and functional changes

We further predicted an increased likelihood of cancer development linked to nine specific mutations, including L381R, A432T, R614G, G799R, R806H, R528C, R528H, V781M, and R671G. These nsSNPs were divided into two groups. The first group, referred to as "covered mutations", included L381R, A432T, R614G, G799R, and R806H. The second group, known as "clustered mutations", consisted of R528C, R528H, V781M, and R671G (**Fig. S3**). These nine nsSNPs were prioritized for further investigation due to their potential cancer association. Additionally, we identified two nsSNPs (Y316C and R643W) as potentially harmful. The results included g-scores, which indicate pathogenicity, and p-values. A g-score above 0.50 suggests a mutation is likely pathogenic. Both Y316C and R643W demonstrated significant pathogenic potential, with g-scores greater than 0.80 and p-values below 0.05, highlighting their importance for further research (**Table 1**).

**Table 1:** Functional and structural modifications of *LIG3* gene predicted by MutPred2.

| Mutation | Actionable/confident hypothesis | g-value | p-value | Probability | Affected PROSITE and ELM motifs |
| --- | --- | --- | --- | --- | --- |
| R528C | Altered DNA binding | 0.919 | 0.34 | 1.2e-03 | ELME000063,<br>ELME000106,<br>ELME000146,<br>ELME000182,<br>ELME000336 |
|  | Altered Ordered interface |  | 0.27 | 0.04 |  |
|  | Altered Metal binding |  | 0.18 | 0.05 |  |
|  | Altered Transmembrane protein |  | 0.16 | 0.01 |  |
|  | Loss of Catalytic site at R528 |  | 0.08 | 0.05 |  |
| R614G | Altered Stability | 0.882 | 0.12 | 0.03 | ELME000102,<br>ELME000108,<br>ELME000146 |
| V781M | Loss of Relative solvent accessibility | 0.824 | 0.26 | 0.03 | None |
|  | Altered Metal binding |  | 0.19 | 0.03 |  |
|  | Loss of Methylation at K778 |  | 0.09 | 0.05 |  |
|  | Altered Stability |  | 0.09 | 0.05 |  |
| R671G | Altered DNA binding | 0.935 | 0.27 | 4.2e-03 | ELME000093,<br>ELME000100,<br>ELME000108,<br>ELME000233,<br>PS00009,<br>PS00333 |
|  | Gain of Strand |  | 0.26 | 0.04 |  |
|  | Gain of Loop |  | 0.26 | 0.05 |  |
|  | Loss of Acetylation at K670 |  | 0.25 | 0.01 |  |
|  | Altered Ordered interface |  | 0.024 | 0.04 |  |
| G799R | Altered Ordered interface | 0.845 | 0.27 | 0.05 | ELME000062 |
|  | Altered Disordered interface |  | 0.27 | 0.05 |  |
|  | Loss of Strand |  | 0.27 | 0.03 |  |
|  | Altered Metal binding |  | 0.22 | 0.02 |  |
|  | Altered Metal binding |  | 0.21 | 0.02 |  |
|  | Altered Transmembrane protein |  | 0.12 | 0.03 |  |
|  | Loss of Proteolytic cleavage at R803 |  | 0.11 | 0.04 |  |
| L381R | Gain of Intrinsic disorder | 0.936 | 0.32 | 0.03 | ELME000012,<br>ELME000231,<br>ELME000313,<br>ELME000333 |
|  | Altered Transmembrane protein |  | 0.20 | 5.7e-03 |  |
|  | Altered Stability |  | 0.10 | 0.04 |  |
| A432T | Gain of Acetylation at K427 | 0.779 | 0.23 | 0.02 |  |
|  | Gain of Allosteric site at K427 |  | 0.22 | 0.03 |  |
|  | Loss of Ubiquitylation at K427 |  | 0.16 | 0.04 |  |
| R806H | Altered Disordered interface | 0.859 | 0.28 | 0.04 | ELME000062 |
|  | Gain of Strand |  | 0.26 | 0.04 |  |
|  | Altered DNA binding |  | 0.22 | 0.01 |  |
|  | Loss of Proteolytic cleavage at R803 |  | 0.11 | 0.04 |  |
| R528H | Altered DNA binding | 0.827 | 0.29 | 3.6e-03 | ELME000063,<br>ELME000106,<br>ELME000146,<br>ELME000182,<br>ELME000336 |
|  | Altered Metal binding |  | 0.19 | 0.05 |  |
|  | Altered Transmembrane protein |  | 0.14 | 0.02 |  |

### 3.4 Estimating the effects of high risk nsSNPs on the structure of protein

The structural effects of high-risk nsSNPs on the LIG3 protein revealed distinct physicochemical changes between the wild-type and mutant amino acids, including variations in size, charge, and hydrophobicity. Among the nine identified nsSNPs, four mutations (V781M, L381R, A432T, and G799R) led to an increase in amino acid size, while five mutations (R528C, R671G, R528H, R614G, and R806H) resulted in size reductions. Eight of these mutations also altered the charges of the amino acids. Furthermore, mutations R528C, R671G, R528H, R614G, and R806H exhibited reduced hydrophobicity compared to their wild-type counterparts, while L381R, A432T, and G799R showed increased hydrophobicity, which could potentially influence hydrophobic interactions within the protein structure (**Table 2**). These results revealed that the mutant-type amino acids diverged markedly from the wild-type proteins (**Table S5**). To further investigate, we generated three-dimensional (3D) models of the nine mutant LIG3 proteins. These 3D models, displayed with ribbon representations (**Fig. S3**), clearly highlight the structural changes induced by the mutations, providing valuable insights into the critical aspects of the LIG3 protein.

**Table 2:** Properties of wild-type and mutant-type amino acids regarding size, charge, and hydrophobicity analyzed by Project HOPE.

| Amino acid change | Wild type amino acid |  |  | Mutant type amino acid |  |  |
| --- | --- | --- | --- | --- | --- | --- |
|  | Size | Charge | Hydrophobicity | Size | Charge | Hydrophobicity |
| R671G | Larger | Positive | Less hydrophobic | Smaller | Neutral | More hydrophobic |
| R528C | Larger | Positive | Less hydrophobic | Smaller | Neutral | More hydrophobic |
| V781M | Smaller | - | - | Larger | - | - |
| R528H | Larger | Positive | Less hydrophobic | Smaller | Neutral | More hydrophobic |
| L381R | Smaller | Neutral | More hydrophobic | Larger | Positive | Less hydrophobic |
| A432T | Smaller | Neutral | More hydrophobic | Larger | Negative | Less hydrophobic |
| G799R | Smaller | Neutral | More hydrophobic | Larger | Positive | Less hydrophobic |
| R614G | Larger | Positive | Less hydrophobic | Smaller | Neutral | More hydrophobic |
| R806H | Larger | Positive | Less hydrophobic | Smaller | Neutral | More hydrophobic |

‘\_’ indicates neutral effect

### 3.5 Functional enrichment and signaling pathways analysis

We assessed the biological characteristics of the *LIG3* gene by functionally annotating its principal targets through gene ontology (GO) enrichment analysis. A total of 23 GO terms were generated, with 9 related to (BP), 7 to cellular components (CC), and 7 to molecular functions (MF). The sizes of the nodes represented the associated target genes, while the color gradient, ranging from green to red, indicated p-values from high to low (**Fig. 1**). The Kyoto Encyclopedia of Genes and Genomes (KEGG) provides a comprehensive pathway database, widely used as a knowledge resource for analyzing biological pathways and cellular activities. Using a p-value threshold of less than 0.05, which was strongly associated with the target genes, KEGG enrichment analysis revealed several enriched pathways related to 11 key targets. The network view of significant KEGG pathways linked to LIG3, highlighting its role in DNA repair and cellular processes (**Fig. 2**). LIG3, as a central node, connects to pathways involved in mitochondrial DNA repair (GO:0043504), DNA ligation (GO:0051103, GO:0006266), and mitochondrial DNA metabolic processes (GO:0032042). It is also associated with V(D)J recombination (GO:0033151), which is crucial for immune diversity. Additionally, phenotypic outcomes, such as embryonic lethality (MP:0011106, MP:0011107), reduced embryo size (MP:0001698), increased mitotic sister chromatid exchange (MP:0003701), and growth retardation (MP:0003984), further highlight LIG3’s critical role in genomic stability and development (**Fig. 2**).

**Fig. 2.**
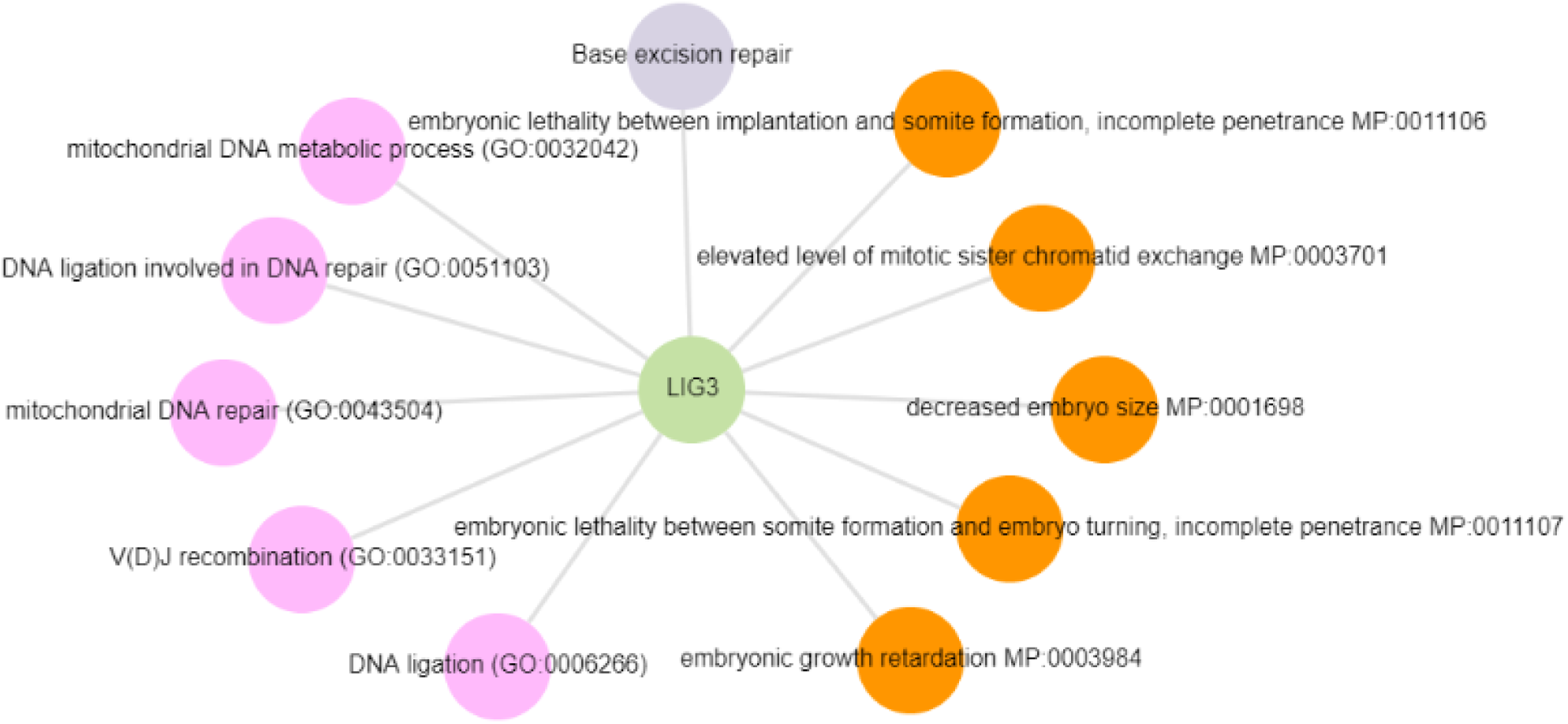
Significant KEGG pathways of *LIG3* were represented in network view. The findings for the pathway term results were sorted based on the combined score (P-value).

### 3.6 Prediction of protein-protein interaction

The PPI network analysis revealed that LIG3 interacts with ten other proteins: APLF, PRKDC, NHEJ1, XRCC6, XRCC4, LIG4, ATM, DCLRE1C, PAXX, and PARP1 (**Fig. S4**). This interaction network, which includes 11 nodes and 55 edges, demonstrates a highly interconnected web, with LIG3 at its center. The network’s high PPI enrichment value of 1.11e-16 and an average node degree of 9.64 suggest significant functional interactions among these proteins, likely involving mutual regulation. Different edge colors were used to visually represent the protein-protein connections (**Fig. S4**).

### 3.7 Superimposition of wild and mutated type proteins

The superimposition of wild-type and mutant LIG3 proteins, performed using the Chimera tool, revealed structural changes induced by mutations (**Fig. 3**). The R528C mutation, where Arginine is replaced by Cystine, led to deviations in the loop region (**Figs. 3A-B**). The R671G mutation, involving the substitution of Arginine with Glycine, resulted in a reduction of side-chain bulkiness, affecting the local structure (**Figs. 3C-D**). The V781M mutation, where Methionine is replaced by Valine, caused alterations in side-chain length and packing (**Figs. 3E-F**). These mutations induced localized structural shifts, which may impact protein stability and ligand interactions. The observed structural deviations, particularly in key residues, suggested significant consequences for *LIG3* functionality and its binding dynamics, emphasizing the relevance of this research.

**Fig. 3.**
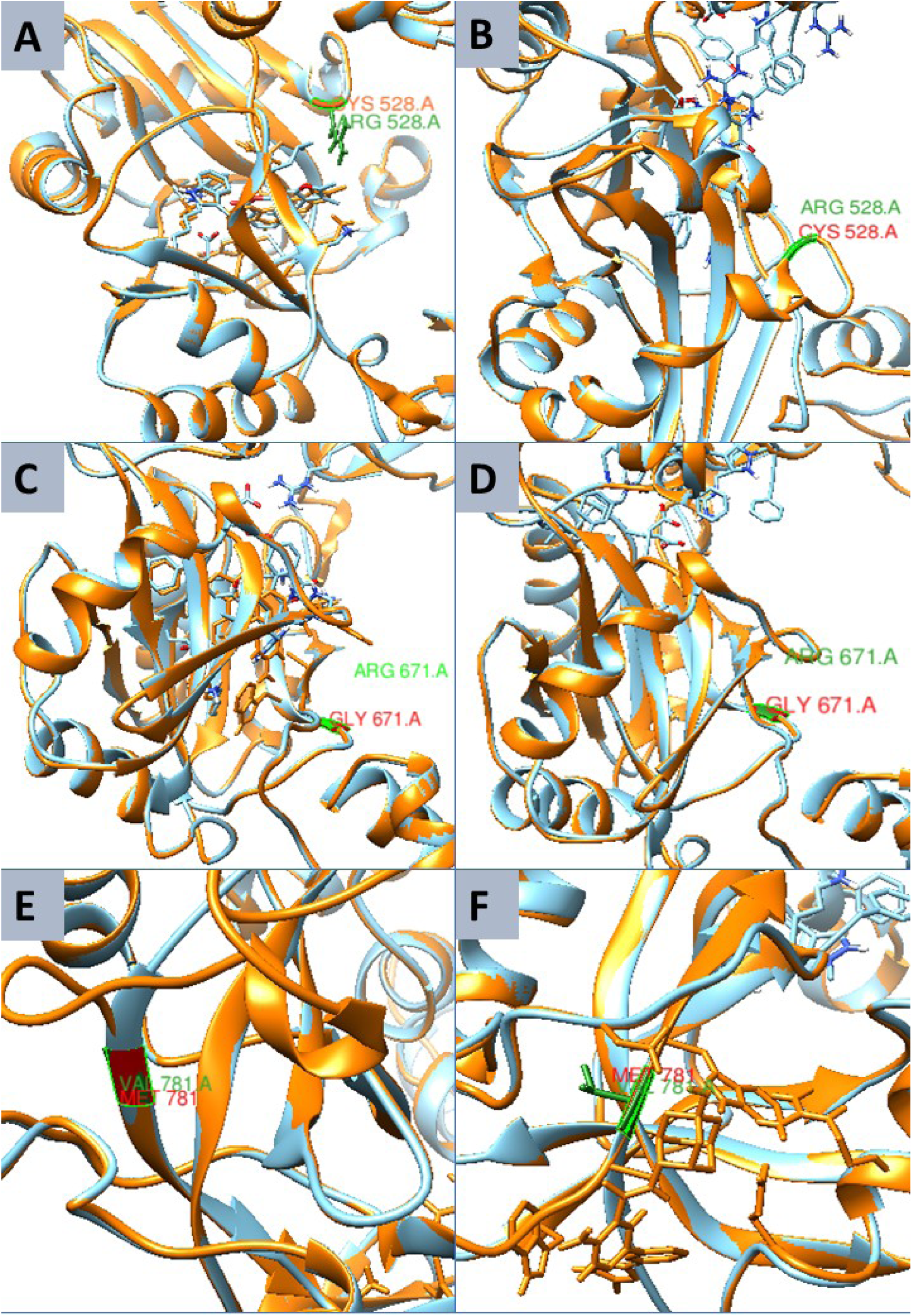
The LIG3 protein in its wild-type form superimposed with three mutant proteins. The structure illustrates the superimposition of the wild-type and mutant LIG3 proteins, with mutations at positions 528 (Arginine to Cysteine) in panel A and B, 671 (Arginine to Glycine) in panel C and D, and 781 (Valine to Methionine) in panel E and F, respectively. Proteins are labeled with colors: orange for wild-type, cyan for mutant, and a yellow box representing the sites of mutations in three mutant proteins relative to one wild-type protein.

### 3.8 Binding interactions

We observed a significant reduction in binding affinity for certain compounds due to the three nsSNPs (**Table S6**). The docked complexes were carefully analyzed for their binding affinity (kcal/mol) and interaction structures (**Fig. 4**). The R528C, V781M, and R671G mutations showed reduced binding affinities for disease-associated compounds, including AHP-MPC (CID: 70687578), DM-BFC (CID: 707801) (**Table 2**), and several others (CID: 116535, CID: 59937, CID: 408383, CID: 676443, CID: 718154, CID: 722325, CID: 609964, and CID: 684700) (**Table S4**). In its native form, the peptide sequence showed substantial hydrogen bonds, yet its binding affinity remains below -10.0 kcal/mol. In contrast, the mutant forms exhibited decreased binding affinities and formed fewer hydrogen bonds with the target molecules.

**Fig. 4.**
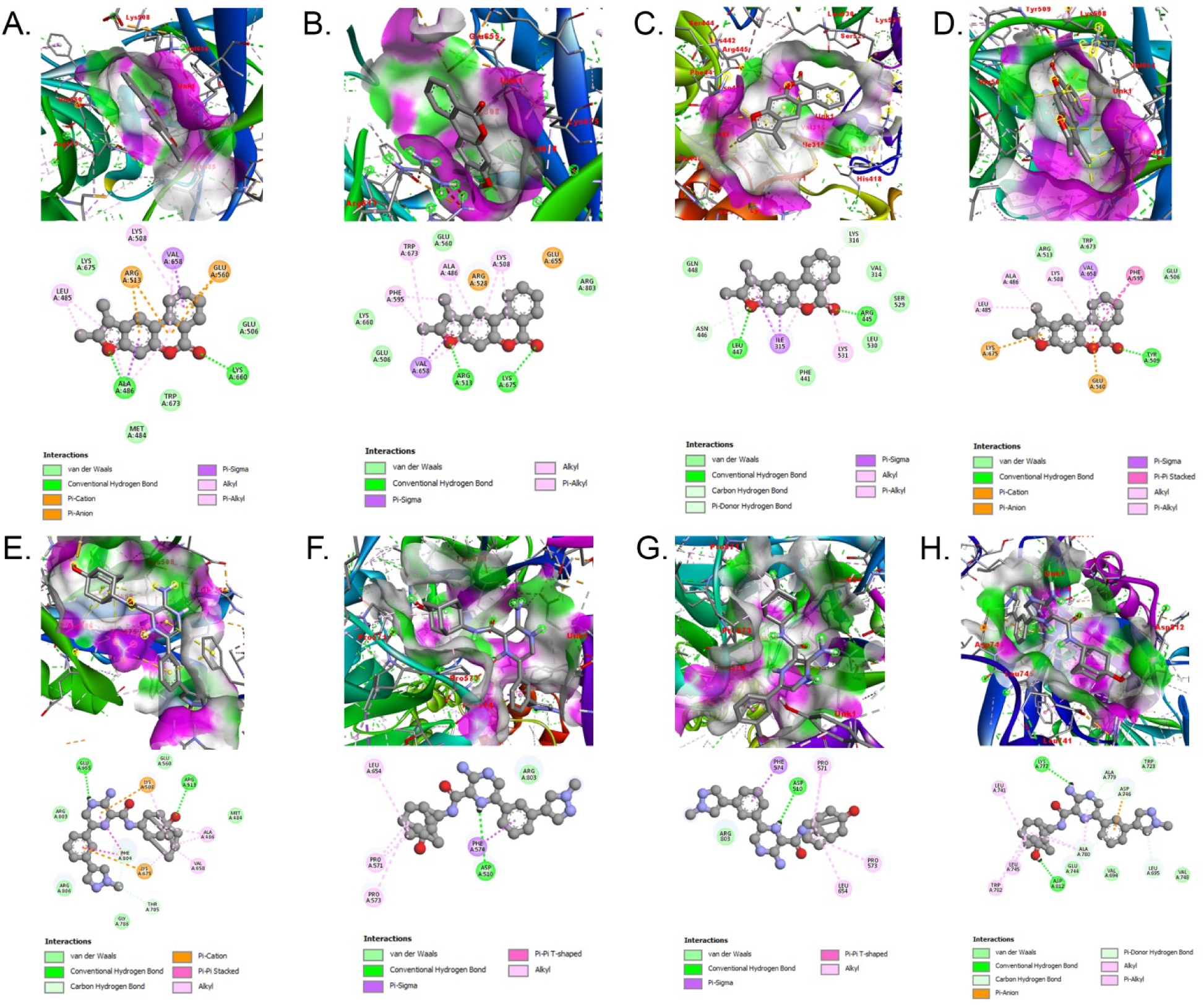
Assessment of binding interactions of wild-type and mutated LIG3 proteins utilizing Autodock Vina. The upper side illustrates the interactions involving (A) R528C with DM-BFC, (B) R671G with DM-BFC, (C) V781M with DM-BFC, (D) the wild-type protein with DM-BFC, (E) the wild-type protein with AHP-MPC, (F) Y316C with AHP-MPC, (G) R643W with AHP-MPC, and (H) the wild-type protein with AHP-MPC.

### 3.9 Pharmacokinetics and toxicity profiles of ten selected compounds

The drug-likeness and ADMET profiles of the AHP-MPC and DM-BFC compounds are presented in **Table 3**. The evaluation of drug-likeness was conducted using Lipinski’s "rule of five," which considers criteria such as molecular weight (MW) < 500 daltons (Da), octanol-water partition coefficient (LOGPo/w) < 5, hydrogen bond donors < 5, and hydrogen bond acceptors < 10. The analysis confirmed that both compounds complied with Lipinski’s guidelines. Furthermore, additional physicochemical properties of the selected compounds such as the number of rotatable bonds, heavy atoms, aromatic heavy atoms, hydrogen bond acceptors, and hydrogen bond donors (detailed in **Table 3**) indicate that these compounds hold promise as safe candidates for therapeutic applications. Protox III was utilized to evaluate the toxicological profiles of the screened compounds. The analysis revealed that both compounds are free from AMES toxicity, hepatotoxicity, and skin sensitization. Furthermore, they demonstrated safety and minimal toxicity in the oral acute toxicity (LD_50_) test conducted on rats. Notably, the favorable ADME profiles and physicochemical properties of these compounds highlight their potential as promising candidates for the development of new medications (**Table 3**).

**Table 3:** Assessment of outlined compounds, focusing on their ADMET (Absorption, Distribution, Metabolism, Excretion, and Toxicity), physicochemical characteristics, lipophilicity, and drug-likeness.

| Properties | Model | DM-BFC | AHP-MPC |
| --- | --- | --- | --- |
| Absorption | Intestinal absorption (human) | 96.726 | 95.895 |
|  | Skin permeability | -2.53 | -2.761 |
|  | Caco2 permeability | 1.409 | 0.525 |
|  | Water solubility | -5.772 | -2.892 |
|  | P-glycoprotein substrate | Yes | No |
|  | P-glycoprotein I inhibitor | No | No |
|  | P-glycoprotein II inhibitor | No | No |
| Distribution | VDss (human) | 0.241 | 0.299 |
|  | Fraction unbound (human) | 0.196 | 0.186 |
|  | BBB permeability | 0.382 | -1.022 |
|  | CNS permeability | -1.306 | -2.564 |
| Metabolism | CYP2D6 substrate | No | No |
|  | CYP3A4 substrate | Yes | Yes |
|  | CYP2D6 inhibitor | No | No |
|  | CYP3A4 inhibitor | No | Yes |
|  | CYP1A2 inhibitor | Yes | Yes |
|  | CYP2C19 inhibitor | Yes | No |
|  | CYP2C9 inhibitor | Yes | No |
|  | Renal OCT2 substrate | No | No |
| Toxicity | AMES toxicity | No | No |
|  | Oral rat acute toxicity (LD<sub>50</sub>) | 0.684 | 3.29 |
|  | Hepatotoxicity | No | No |
|  | Max. tolerated dose (human) | 0.124 | 0.438 |
|  | hERG I inhibitor | No | No |
|  | hERG II inhibitor | Yes | No |
|  | Oral rat chronic toxicity<br>(LOAEL) | 0.684 | 14.01 |
|  | Skin sensitization | No | No |
| Physicochemical | Molecular weight | 264.28 g/mol | 444.53 g/mol |
|  | LogP | 4.30924 | 2.7957 |
|  | No. H-bond acceptors | 3 | 5 |
|  | No. H-bond donors | 0 | 3 |
|  | Molar refractivity | 76.69 | 125.47 |
|  | TPSA | 43.35 Å<sup>2</sup> | 118.95 Å<sup>2</sup> |
| Lipophilicity | Consensus Log $P_{o/w}$ | 3.88 | 2.31 |
| Water Solubility | Log $S$ (ESOL) | -4.75 | -4.12 |
|  | Solubility class | Moderately<br>soluble | Moderately<br>soluble |
| Drug Likeness | Lipinski violation | Yes: 0 violation | Yes: 0 violation |
|  | Bioavailability score | 0.55 | 0.55 |
| Medicinal Chemistry | PAINS | 0 alert | 0 alert |

Here: BBB- blood brain barrier, VDss: volume of distribution at steady state, TPSA: topological polar surface area, PAINS: pan-assay interference compounds.

### 3.10 Molecular dynamics simulation

The study evaluated the interaction of AHP-MPC and DM-BFC compounds with three mutant forms of LIG3 protein namely R528C, V781M, and R671G compared to the wild-type LIG3 protein using 100 ns molecular dynamics simulations (MDS). Results indicated distinct binding modes of the ligands with both wild-type and mutant LIG3 proteins. RMSD analysis showed varying stability profiles among the proteins in the presence of the ligands (**Figs. 5A-B**). AHP-MPC maintained stable conformations with the wild-type and R528C mutant (RMSD values ∼4– 6 Å), whereas R671G and V781M mutants exhibited instability. DM-BFC improved stability across all systems (RMSD values ∼3–6 Å), especially with the wild-type and R528C mutant. These findings suggest that the wild-type and R528C mutant are promising targets due to their stable interactions with both ligands, while R671G and V781M mutations induce structural instability, particularly with AHP-MPC, emphasizing mutation-specific and ligand-dependent effects on LIG3 protein dynamics (**Figs. 5A-B**). RMSF analysis further highlighted structural differences among the wild-type LIG3 protein and its mutants (R528C, R671G, and V781M) in the context of their interactions with AHP-MPC and DM-BFC (**Figs. 5C-D**). The wild-type LIG3 complexed with DM-BFC exhibited the least flexibility (lowest RMSF values), indicating a stable structure. In contrast, R671G and V781M mutants showed increased flexibility, particularly in residues 300– 400 and the C-terminal region beyond residue 500. Notably, the C-terminal region displayed significant fluctuations, especially in the V781M–AHP-MPC complexes. Interestingly, the R528C mutant maintained lower RMSF values comparable to the wild-type in both ligand conditions, underscoring its stability. These results suggest that DM-BFC enhances stability, particularly for the wild-type and R528C mutant proteins, supporting their potential therapeutic applications (**Figs. 5C-D**).

**Fig. 5.**
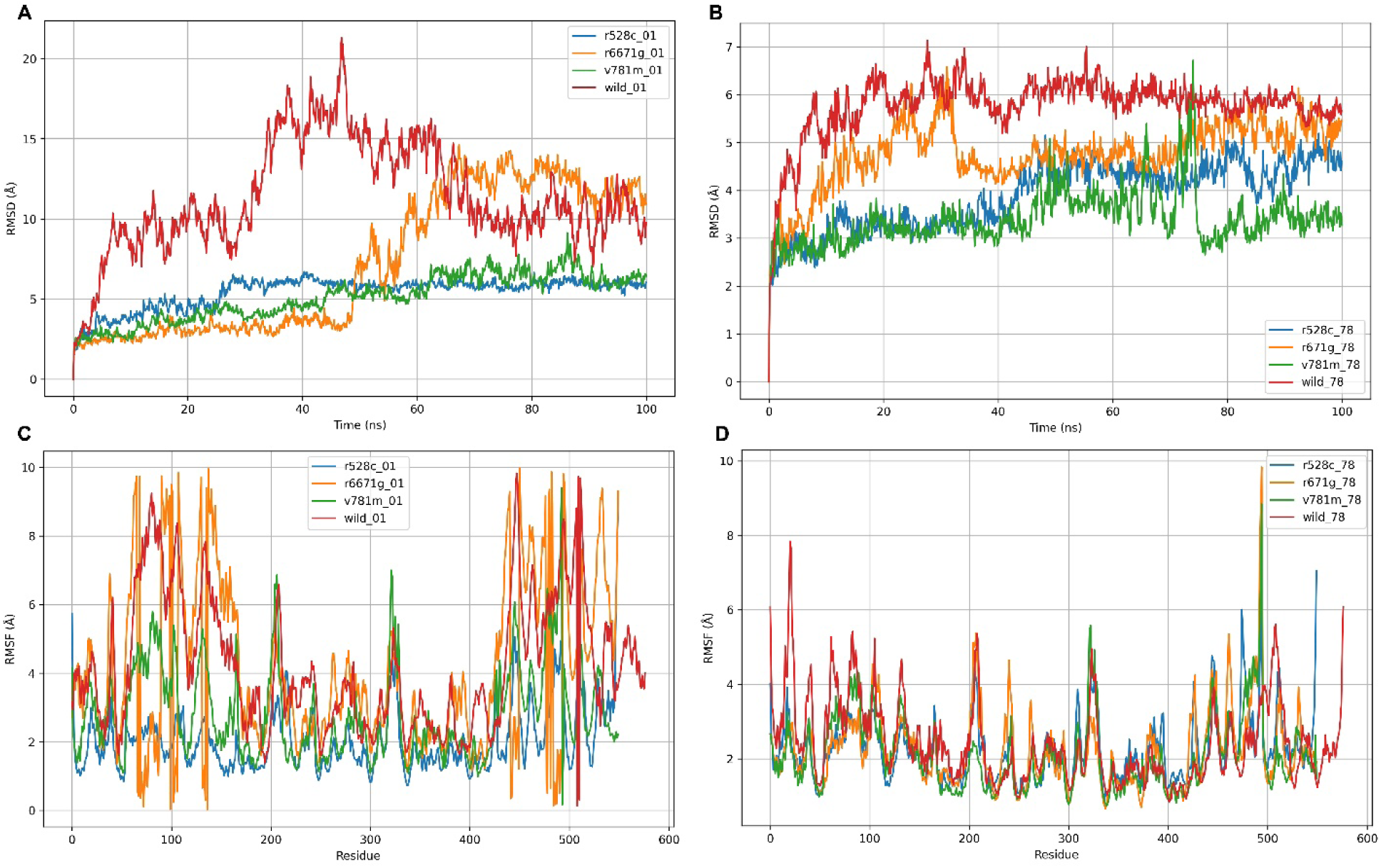
The RMSD values for the wild-type LIG3 protein and three mutant-type LIG3 proteins (R528C, R671G, and V781M) were analyzed in incorporation with the two ligands: DM-BFC (Panel A) and AHP-MPC (Panel B). The root mean square fluctuations (RMSF) values of the wild-type LIG3 protein, three mutant-type LIG3 proteins (R528C, R671G, and V781M), and the two ligands (DM-BFC and AHP-MPC) are utilized to evaluate the structural changes of proteins.

The rGyr analysis revealed the compactness and structural integrity of the wild-type LIG3 protein and its mutant variants with the presence of DM-BFC and AHP-MPC (**Fig. 6A**). Regarding DM-BFC, the wild-type LIG3 protein demonstrated with the highest rGyr values (∼32–36 Å), indicating considerable structural expansion, especially in the early phase. Conversely, the R528C and V781M mutants exhibited persistently lower rGyr values (∼26–28 Å), signifying a more compact and stable conformation. The R671G mutant exhibited moderate rGyr values (about 28– 32 Å). For AHP-MPC, all variants exhibited reduced rGyr values with negligible variations (∼26.5–28.5 Å), signifying improved stability (**Fig. 6B**). Notably, The R528C mutant demonstrated the lowest rGyr values across both ligand conditions, suggesting enhanced compactness and structural integrity. These findings underscore the importance of our research in understanding the structural dynamics of LIG3 protein variants. The SASA analysis revealed the exposure of the wild-type LIG3 protein and its mutants to solvents in the presence of DM-BFC and AHP-MPC. For DM-BFC, the wild-type protein showed the highest SASA values (∼30,000– 31,500 Å²), indicating greater solvent exposure and a relatively expanded structure. In contrast, the R528C and V781M mutants exhibited significantly lower SASA values (∼26,000–28,000 Å²), suggesting reduced solvent accessibility and a more compact structure. The R671G mutant maintained intermediate SASA values (∼28,000–29,000 Å²) (**Fig. 6C**). For AHP-MPC, all variants displayed reduced SASA values compared to DM-BFC. The wild-type protein retained higher SASA values (∼29,000 Å²), while the R528C mutant consistently exhibited the lowest solvent exposure, indicating superior structural compactness (**Fig. 6D**). The PSA and MolSA analyses demonstrated substantial differences between the wild-type LIG3 protein and its mutant variants in the presence of DM-BFC and AHP-MPC (**Fig. 7**). For PSA, the wild-type protein exhibited the highest values with DM-BFC (∼15,000 Å²), suggesting an increased solvent-accessible surface area of polar regions, whereas the R528C and V781M mutants displayed lower values (∼13,500– 14,000 Å²), indicative of a more compact conformation. The R671G mutant showed intermediate PSA value (∼14,200 Å²) (**Fig. 7A**). Upon interaction with AHP-MPC, PSA values decreased for all variants (∼13,200–14,000 Å²), implying ligand-induced stabilization (**Fig. 7B**). Simultaneously, MolSA analysis revealed that the wild-type protein had the highest MolSA value (∼27,000 Å²), particularly with DM-BFC, while the R528C mutant displayed the lowest value (∼24,000 Å²) (**Fig. 7C).** The MolSA values further diminished in the presence of AHP-MPC, reinforcing the compact structural conformation (**Fig. 7D**).

**Fig. 6.**
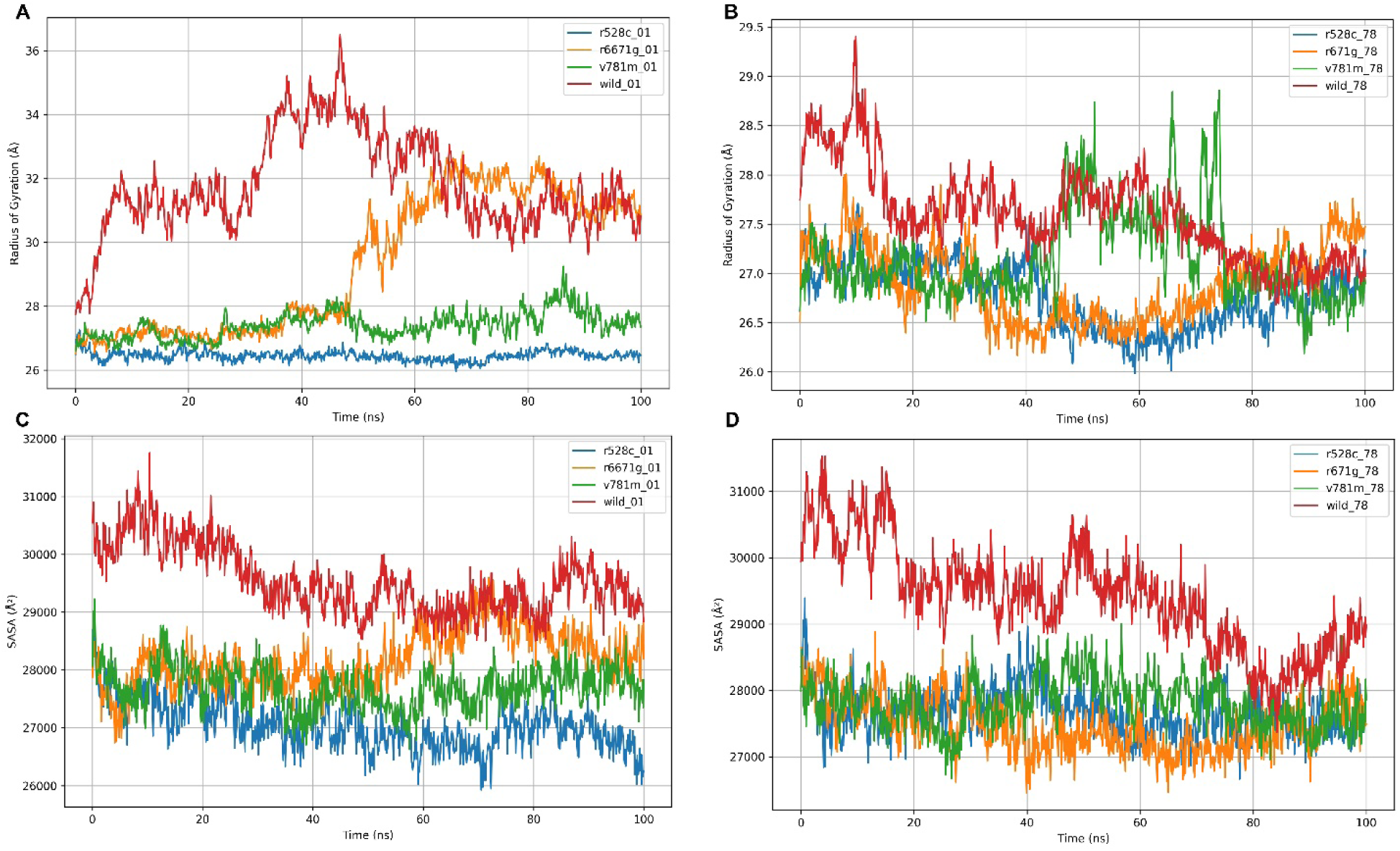
The radius of gyration (rGyr) values for the wild-type LIG3 protein and three mutant-type LIG3 proteins (R528C, R671G, and V781M) were confronted with the two ligands (DM-BFC and AHP-MPC). The wild-type LIG3 protein and three mutant-type LIG3 proteins (R528C, R671G, and V781M) were assessed utilizing solvent accessible surface area (SASA) values considering interacting with two ligands (DM-BFCand AHP-MPC).

**Fig. 7.**
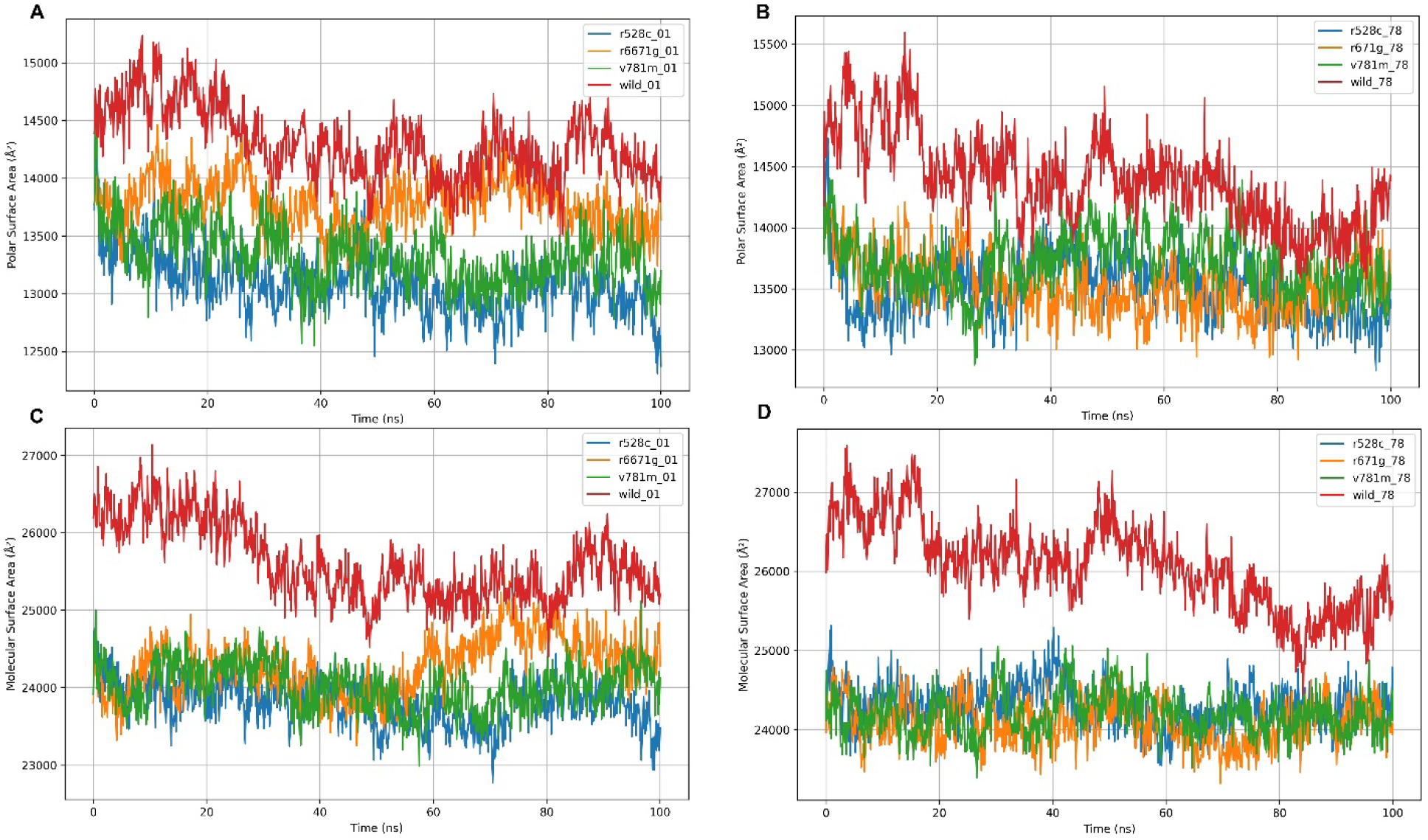
Polar surface area (Panel A and B) and Molecular surface area (Panel C and D) values of the wild-type LIG3 protein and three mutant-type LIG3 proteins (R528C, R671G and V781M) were interacting with the two ligands (DM-BFCand AHP-MPC).

### 3.11 Association of *LIG3* gene in various cancer

We further explored the association between the *LIG3* gene and the survival rate for patients with breast cancer, bladder cancer, AML, and hepatocellular carcinoma. The Kaplan-Meier analysis demonstrated a significant correlation between high *LIG3* expression and improved survival in breast cancer (HR: 0.81, p = 0.00093), and AML (HR: 0.67, p = 3e-05), respectively. However, no significant association was observed in bladder cancer (HR: 0.93, p = 0.16) or hepatocellular carcinoma (HR: 0.93, p = 0.32), and it demonstrated reduced survival rates for these cancer types (**Fig. 8**). These findings suggest that LIG3 may serve as a prognostic biomarker in breast cancer and AML but not in bladder or liver cancers, underscoring its potential cancer-type-specific relevance in survival outcomes.

**Fig. 8.**
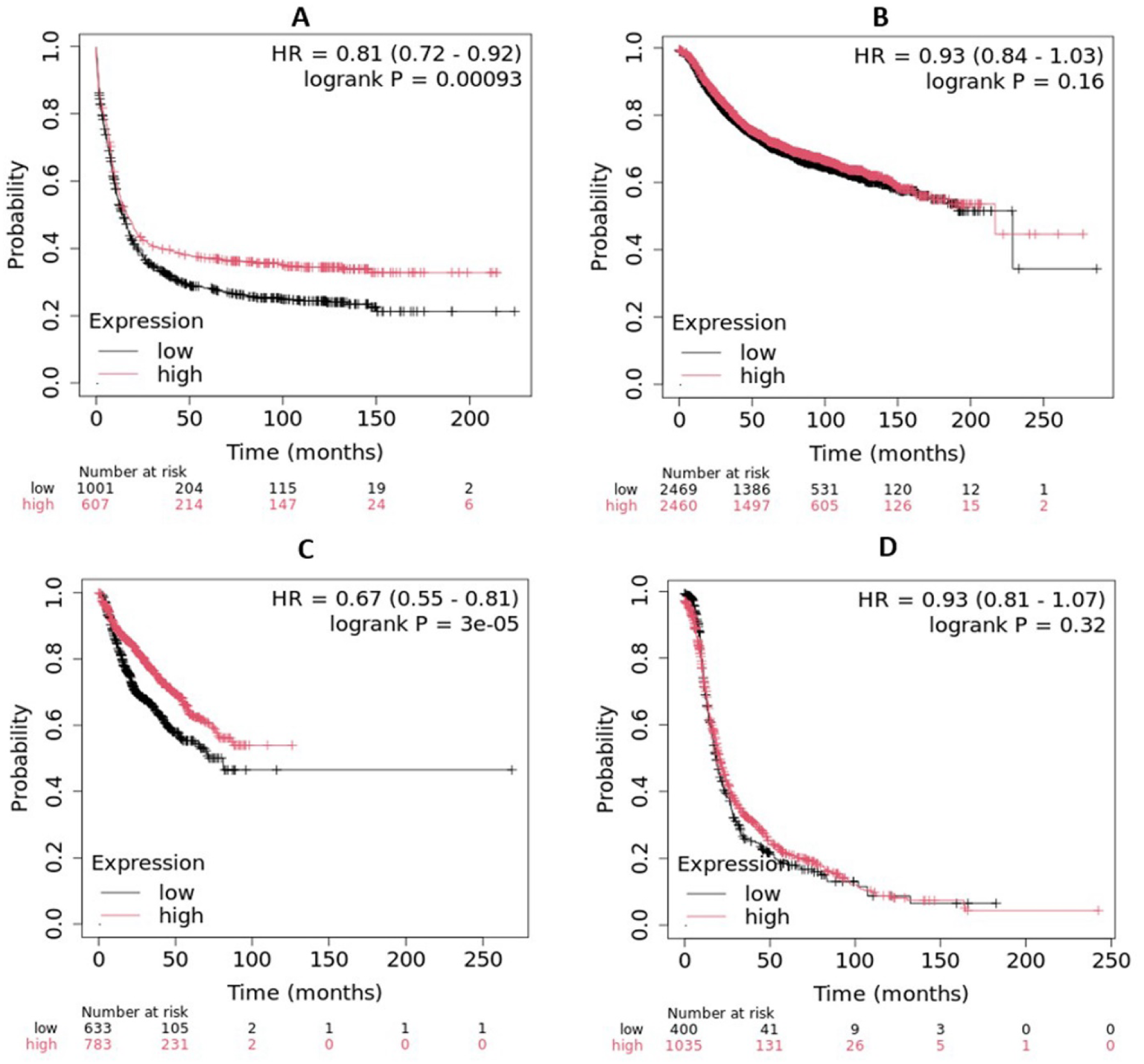
Level of *LIG3* gene expression and survival rates in patients with different forms of cancer (A: breast cancer, B: bladder cancer, C: AML and D: hepatocellular carcinoma) utilizing microarray data by Kaplan-Meirer Plotter. "P" denotes p-values, while "HR" stands for hazard ratio.

## 4. Discussion

This study intends to uncover the potential influence of genetic variants (nsSNPs) on gene function, particularly their effects on protein structure, function, and their relevance to AML through an analysis of the *LIG3* gene. We identified 12,191 nsSNPs in the *LIG3* gene, with 902 (7.4%) being missense variants. Among these, 18 were classified as detrimental due to their potential to disrupt protein structure or function. This finding aligns with previous studies that highlight the importance of missense mutations in altering protein function, particularly in DNA repair genes like *LIG3* [59, 60]. The identification of 132 somatic missense variants further underscores the relevance of *LIG3* in cancer biology, as somatic mutations are often drivers of oncogenesis [61]. We further predicted that 12 out of 18 detrimental nsSNPs significantly reduced protein stability, while the other SNPs demonstrated elevated stability. We excluded mutations that stabilize the protein because this wasn’t the intended purpose of our research, as changes in protein stability affect its structural conformation and functions [62]. Protein stability is crucial for maintaining functional integrity, and destabilizing mutations can lead to loss of function or misfolding, which is often associated with disease [63]. The identification of potential methylation and phosphorylation sites (224R, 529S, and 666Y) further highlights the role of post-translational modifications in regulating *LIG3* gene activity. These findings align with previous research indicating that phosphorylation and methylation can regulate the activity of DNA repair proteins, highlighting their crucial role in connecting genetic variations to phenotypic outcomes [59, 60, 64]. Nine nsSNPs were linked to an increased risk of AML, showing significant potential for oncogenic transformation. The amino acid size was found to be increased in the wild-type variant due to mutations V781M, L381R, A432T, and G799R and decreased due to mutations R528C, R671G, R528H, R614G, and R806H. Changes in protein size due to mutations can disrupt the folding and spatial arrangement of the protein, altering its stability and functionality [65]. The classification into "covered" and "clustered" mutations provides a framework for understanding their structural and functional impacts. Notably, Y316C and R643W were identified as highly pathogenic, with g-scores > 0.80, suggesting their potential as biomarkers for cancer risk assessment. This aligns with studies that have identified specific nsSNPs in DNA repair genes as predictive markers for cancer susceptibility [66, 67].

The study revealed significant physicochemical changes in the LIG3 protein due to high-risk nsSNPs, including alterations in size, charge, and hydrophobicity. These changes can disrupt PPI and ligand binding, which are critical for LIG3’s role in DNA repair. Charge modifications may affect active sites along with PPI, resulting in functional impairments, such as ineffective DNA repair, which may play a role in diseases like cancer [68, 69]. The 3D models of mutant proteins provided visual evidence of structural deviations, supporting the hypothesis that these mutations impair protein function. Similar structural analyses have been used to elucidate the impact of nsSNPs in other DNA repair proteins, such as BRCA1 [70]. GO and KEGG pathway analyses highlighted the involvement of *LIG3* gene in DNA repair, mitochondrial DNA metabolism, and V(D)J recombination. The interaction of the LIG3 protein with other ligase proteins, such as LIG1 and LIG4, suggests a multifaceted role in DNA repair pathways that may influence the formation and progression of AML [10]. Furthermore, GO enrichment analysis of the *LIG3* gene provided valuable insights into its function in DNA repair and genomic integrity by identifying over-represented GO terms associated with LIG3 [71]. These findings are consistent with LIG3’s known role in maintaining genomic stability [72]. The PPI network analysis further revealed the interactions of LIG3 protein with key DNA repair proteins, including XRCC4, PARP1, and ATM. These interactions are critical for NHEJ and base excision repair (BER), pathways essential for maintaining genomic integrity [73]. The high PPI enrichment value (1.11e-16) suggests that LIG3 is a central player in these pathways, further emphasizing its importance in cancer biology [74]. The superimposition of wild-type and mutant LIG3 proteins revealed significant structural deviations, particularly in loop regions and side-chain packing. These changes can affect protein stability and ligand binding, as seen in other DNA repair proteins like XRCC1 [75]. The observed structural shifts provide a mechanistic basis for the functional impairments associated with these mutations. We found that mutations like R528C, V781M, and R671G reduced binding affinity for disease-associated compounds. This is consistent with previous research showing that nsSNPs can disrupt ligand binding, leading to functional deficits [76]. The reduced hydrogen bonding in mutant forms further supports the idea that these mutations impair protein-ligand interactions. The drug-likeness and ADMET profiles of AHP-MPC and DM-BFC suggested that these compounds are promising candidates (molecular weights <500 g/mol) for therapeutic development. Their compliance with Lipinski’s rule of five and favorable toxicity profiles align with criteria for drug development [56]. These findings are significant for developing targeted therapies in cancers linked to *LIG3* dysfunction, suggesting that the identified compounds could serve as promising candidates for future in vivo drug evaluations for AML.

Molecular dynamics (MD) simulation proved to be a robust computational approach in drug discovery, offering detailed insights into the conformational dynamics, stability, and interaction mechanisms of protein-ligand complexes [77, 78]. The MDS results revealed that the R528C mutant maintained stability comparable to the wild-type protein, while R671G and V781M exhibited structural instability. These findings are consistent with studies showing that specific mutations can induce conformational changes that affect protein dynamics [79]. The stability of the R528C mutant suggests it may retain partial function, making it a potential target for therapeutic intervention. The Kaplan-Meier analysis demonstrated that high *LIG3* expression correlates with improved survival in breast cancer and AML but not in bladder or liver cancers. This cancer-type-specific association highlights the complex role of LIG3 in different malignancies. Our findings highlighted the cancer-type-specific relevance of *LIG3* expression in survival outcomes, warranting further investigation into its functional mechanisms. Similar findings have been reported for other DNA repair genes, such as BRCA1 and BRCA2, which show tissue-specific effects in cancer prognosis [70, 78].

## Conclusion

AML is a genetically heterogeneous malignancy driven by multiple mutations that influence its initiation, progression, and therapeutic response. This study presents a comprehensive analysis of the *LIG3* gene, a key component of the NHEJ pathway, which plays a critical role in DNA repair, particularly during DNA DSBs repair. Through the analysis of 132 missense SNPs, we identified 12 destabilizing mutations, nine of which were associated with cancer, suggesting their potential pathogenic impact on protein structure and function. Molecular docking studies identified two promising ligands, DM-BFC and AHP-MPC, exhibiting high binding affinity for both mutant and wild-type LIG3 proteins, indicating their potential as therapeutic agents. Pharmacokinetic and dynamics simulation further supported the suitability of these compounds for future preclinical evaluation. These findings highlight the critical role of *LIG3* gene in maintaining genomic integrity and suggest its potential as a therapeutic target for AML. However, further experimental validation is required to confirm these in-silico predictions, elucidate the molecular mechanisms underlying the identified mutations, and evaluate the clinical applicability of the proposed ligands. This study provides a foundation for the development of targeted therapeutics and personalized treatment strategies for AML and other diseases associated with *LIG3* gene dysfunction.

## Data availability

The article contains the data utilized to support the results of the *in-silico* study.

## Author contribution statement

MAH, UMSJ, MNU, TI and MNH: conceptualized and designed the study; MAH, KHA, AR, SA, MMU, MFA, MAB MJI, MK and SH performed experiment, acquired and analyzed data, interpreted results and drafted the original manuscript; MNU, TI and MNH critically reviewed and edited the manuscript. Finally, all authors read, edited and approved the manuscript.

## Ethical statement

This *in-silico* computational study did not involve human subjects and therefore did not require ethical approval. Furthermore, the authors declare that this manuscript, submitted to PLOS ONE, has been prepared with full adherence to responsible research practices and in accordance with the guidelines of publication ethics.

## Consent for Publication

Not applicable.

